# Homologous recombination between tandem paralogues drives evolution of a subset of Type VII secretion system immunity genes in firmicute bacteria

**DOI:** 10.1101/2022.01.07.475358

**Authors:** Stephen R. Garrett, Giuseppina Mariano, Jo Dicks, Tracy Palmer

## Abstract

The Type VII secretion system (T7SS) is found in many Gram-positive firmicutes and secretes protein toxins that mediate bacterial antagonism. Two T7SS toxins have been identified in *Staphylococcus aureus*, EsaD a nuclease toxin that is counteracted by the EsaG immunity protein, and TspA, which has membrane depolarising activity and is neutralised by TsaI. Both toxins are polymorphic, and strings of non-identical *esaG* and *tsaI* immunity genes are encoded in all *S. aureus* strains. To investigate the evolution of *esaG* repertoires, we analysed the sequences of the tandem *esaG* genes and their encoded proteins. We identified three blocks of high sequence similarity shared by all *esaG* genes and identified evidence of extensive recombination events between *esaG* paralogues facilitated through these conserved sequence blocks. Recombination between these blocks accounts for loss and expansion of *esaG* genes in *S. aureus* genomes and we identified evidence of such events during evolution of strains in clonal complex 8. TipC, an immunity protein for the TelC lipid II phosphatase toxin secreted by the streptococcal T7SS, is also encoded by multiple gene paralogues. Two blocks of high sequence similarity locate to the 5’ and 3’ end of *tipC* genes, and we found strong evidence for recombination between *tipC* paralogues encoded by *Streptococcus mitis* BCC08. By contrast, we found only a single homology block across *tsaI* genes, and little evidence for intergenic recombination within this gene family. We conclude that homologous recombination is one of the drivers for the evolution of T7SS immunity gene clusters.

**DATA SUMMARY:** All sequence data for strains used in this study are available on NCBI under BioProject PRJNA789916. Sequences from the NCTC3000 project are available on NCBI under BioProject PRJEB6403. Supplementary data 2 and all custom scripts are available on Github: https://github.com/GM110Z/Garret-et-al.-recombination-paper.

**IMPACT STATEMENT:** The type VII secretion system (T7SS) in firmicutes secretes polymorphic toxins that target other bacteria. To protect from the action of these toxins, bacteria carry multiple paralogous copies of immunity protein-encoding genes that are sequence-related but non-identical. To date, little is known about how T7 immunity gene families evolve. In this study we analysed a cluster of EsaG-encoding genes in *Staphylococcus aureus* which are found at the *ess/T7* secretion locus and provide immunity against the T7 secreted nuclease toxin, EsaD. We identified three homology blocks covering *esaG* genes and their downstream intergenic regions, which are separated by two variable regions. We have shown that recombination can occur between these homology blocks, leading to loss or expansion of *esaG* genes at this locus. Using a historical dataset of closely related *S. aureus* strains from clonal complex 8, we identified several independent recombination events leading to changes in the *esaG* repertoire. We further showed that similar events are observed for an immunity protein encoded by Group B *Streptococcus* spp. suggesting that recombination plays a broader role in the evolution of T7SS immunity-encoding genes. We speculate that gain and loss of T7 immunity genes is weighed in response to environmental pressure and metabolic burden.

## INTRODUCTION

The type VII protein secretion system (T7SS) is found in many Gram-positive bacteria. Following its discovery in pathogenic Mycobacteria, it has since been described in a range of other actinobacteria and in firmicutes (1-5). Cryo-electron microscopy studies have shown that the ESX-5 T7SS from Mycobacteria exists as a 2.3 MDa membrane complex, with a central ATPase, EccC, forming a hexameric pore (6, 7). The firmicutes T7SS is only distantly related to the actinobacterial T7SSa system and has been termed T7SSb. The hexameric ATPase is the only common component found across all T7SSs and is designated EssC in the T7SSb systems (8, 9).

While the T7SSa is heavily linked with Mycobacterial virulence, there is growing evidence that the T7SSb plays an important role in bacterial antagonism (10, 11). In *Streptococcus intermedius*, three T7SSb-secreted antibacterial effectors have been identified, including TelB, an NADase, and TelC a lipid II phosphatase (12). The *Enterococcus faecalis* T7SSb also mediates contact-dependent inhibition of some firmicute bacteria, and a bioinformatic analysis in *Listeria monocytogenes* has identified over 40 potential antibacterial substrates of the T7SS (13, 14).

A role for the T7SSb in interbacterial competition was first characterised in the opportunistic pathogen *Staphylococcus aureus* (15). The T7SS-encoding locus in this organism is highly variable. Sequence divergence initiates towards the 3’ end of *essC*, with *essC* sequences falling into one of four variants, termed *ess1* – *ess4* (16). Downstream of each *essC* subtype is a cluster of variant-specific genes. In *essC1* variant strains one of these genes, *esaD*, encodes a secreted nuclease toxin with antibacterial activity. Protection from the toxic activity of EsaD is mediated by EsaG, which is encoded immediately adjacent to *esaD* in *essC1* strains. EsaG inactivates EsaD by forming a tight complex with the EsaD nuclease domain (15). While all *essC1* strains encode *esaD*, the toxin nuclease domain is polymorphic, and these strains also encode additional copies of *esaG* genes in a highly variable immunity gene island located at the 3’ end of the *ess*/*T7SS* locus (10, 15, 16). These additional *esaG* copies are genetically diverse but share some core regions of similarity within the encoded amino acid sequences (17, 18). Strings of up to ten non-identical *esaG* genes are also found in a similar genomic location in *essC2, essC3* and *essC4* strains (10 and our Supplementary dataset 1).

TspA is a second antibacterial toxin secreted by the *S. aureus* T7SS. TspA has a C-terminal membrane-depolarising domain, and immunity from intoxication is provided by the membrane-bound TsaI protein (19). The *tspA-tsaI* locus is encoded distantly from the T7 gene cluster and is found across all four *essC* variant strains. Similar to EsaD, the TspA toxin domain is polymorphic, and all strains encode clusters of TsaI variants (between two and sixteen copies) directly downstream of *tspA* (19).

It is common for bacteria to encode repertoires of immunity proteins for protection against polymorphic effector proteins. For example, bacteria in the gut accumulate immunity genes in genomic islands that provide protection against type VI secretion system effectors (20). This includes orphan immunity genes in strains that do not encode the cognate effector protein, as we observe for *esaG* genes in *S. aureus essC2, essC3* and *essC4* strains that do not contain *esaD*. While it is common for immunity islands to carry many predicted immunity genes, it is less common to see so many homologues of the same immunity gene clustered together. At present, little is known about the origin of the T7SS immunity repertoires.

In this study we used gene phylogeny and cluster analyses to identify the processes that drive the accumulation of T7SS immunity genes. Our findings indicate that intergenic recombination is a major factor in the expansion and deletion of *esaG* genes, and that this is at play during the evolution of *S. aureus* strains. We also noted that a similar process drives the evolution of *tipC* genes, which encode Streptococcal TelC immunity proteins (12). By contrast, there is limited evidence of intergenic recombination within *tsaI* clusters, and it does not appear to be the main mechanism of evolution for this gene family.

## METHODS

### Strains and genome sequencing

Strain RN6390 was obtained from Professor Jan Maarten van Dijl (University of Groningen, NL). Enhanced genome sequencing was carried out by MicrobesNG (Birmingham, UK). Sequence data for RN6390 can be found under BioProject: PRJNA789916.

### Gene and protein alignment

Nucleotide sequences were obtained from NCBI and aligned with MAFFT v7.489 (21). Amino acid sequences were obtained from NCBI and aligned using MUSCLE v3.8.1551 (22). To construct similarity plots for both nucleotide and amino acid sequences, Plotcon (https://www.bioinformatics.nl/cgi-bin/emboss/plotcon) was executed using aligned sequences, with a window size of 5. For *esaG4*, the full pseudogene was used for the nucleotide alignment. For the alignment of the EsaG4 amino acid sequence, the two predicted open reading frames (ORFs) were used. For Plotcon analysis on large input sequences, alignments were manually curated to remove partial sequences and pseudogenes.

### Recombination Prediction

Aligned nucleotide sequences were opened in the RDP4 software (23) and Run All selected. TrimAl v1.2 (24) was used to remove the unaligned 5’ region of *tsaI* genes before RDP4 analysis was carried out.

### Gene phylogeny

Maximum likelihood trees for nucleotide sequences were built with IQTREE v 2.1.4 (25), with 1000 ultrafast bootstraps. Trees were visualised and annotated using iTOL (26).

### Comparison of genetic loci and gene cluster analysis

T7SS-encoding loci from each *essC* variant were subjected to pairwise comparisons with the RN6390 *T7SS* locus using BLAST (27). The report produced by BLAST was used as a comparison file in genoPlotR package (28) within RStudio (29) to plot regions of similarity between the two loci. FlaGs (30) was used to assess the variation in number of *tsaI* repeats across *S. aureus* strains. Examples were selected to represent diversity at this locus. A bash script was written to extract *ess/T7* loci from 133 *Staphylococcus aureus* strains whole-genome sequenced within the NCTC3000 project (PRJEB6403). Nucleotide profile Hidden Markov Models for genes *essC, esaD, esaG* (generated using all variants found within NCTC8325) and *focA* were used to search with HMMER (31) for the presence and copy number of key genes and to extract locus co-ordinates. In particular, the presence of *esaD* was used to identify *essC1* variant strains and the number of *esaG* copies was recorded. Custom bash scripts were used to extract *esaG* and *DUF5079* genes from the *ess/T7* loci of NCTC CC8 strains and alignments carried out as described. Easyfig was used to analyse the *ess/T7* clusters of NCTC CC8 strains by pairwise comparison and subsequently indicate regions of high sequence similarity between each. These were ordered based on when they were accessioned within the NCTC culture collection.

## RESULTS

### RN6390 encodes twelve *esaG* copies

Much of our previous work on the *S. aureus* T7SS has used strain RN6390, which is a derivative of the parental strain NCTC8325. Two genome sequences are available for NCTC8325, NCTC8325-Oklahoma (GenBank accession number CP000253) and NCTC8325-Sanger (GenBank accession number LS483365). The sequences are closely related but differ primarily by the number of *esaG* genes at the T7 locus, with six in the Oklahoma sequence and 12 in the Sanger sequence. To determine the number of *esaG* copies present in RN6390, we undertook whole genome sequencing (deposited under GenBank accession number CP90001.1), finding 12 *esaG* genes, of identical sequence to those in the NCTC8325-Sanger sequence. We noted that the DNA sequence is highly repetitive in this region, which may have confounded sequence assembly for the Oklahoma accession. Nonetheless our sequence analysis shows that our model strain, RN6390, harbours 12 copies of *esaG*.

### Intergenic recombination is a driving force in the evolution of *esaG* genes

As the sequence at the *esaG* locus of RN6390 is repetitive, we examined this region of the chromosome in more detail. Of the 12 homologues of *esaG*, encoded in this region, the first copy is found directly downstream of *esaD* and encodes the cognate immunity protein for the nuclease toxin (Fig 1a; 15). We have named this *esaG1*. This is followed by three genes encoding hypothetical proteins (SAOUHSC_00270 _00271 and _00272), before a stretch of four further *esaG* homologues (*esaG2* - *esaG5*). This is followed by a further homologue of *SAOUHSC_00270* (annotated as *SAOUHSC*_*00270b* on Fig 1a, sharing 96.49% identity with *SAOUHSC_00270*) and then an additional seven *esaG* homologues (*esaG6* – *esaG12*; Fig 1a). Note that while *esaG4* is annotated as a pseudogene, due to a short additional stretch of nucleotides close to the centre of the gene introducing a premature stop and subsequent start codon, it actually encodes two smaller ORFs (Fig 1a).

**Figure 1.**
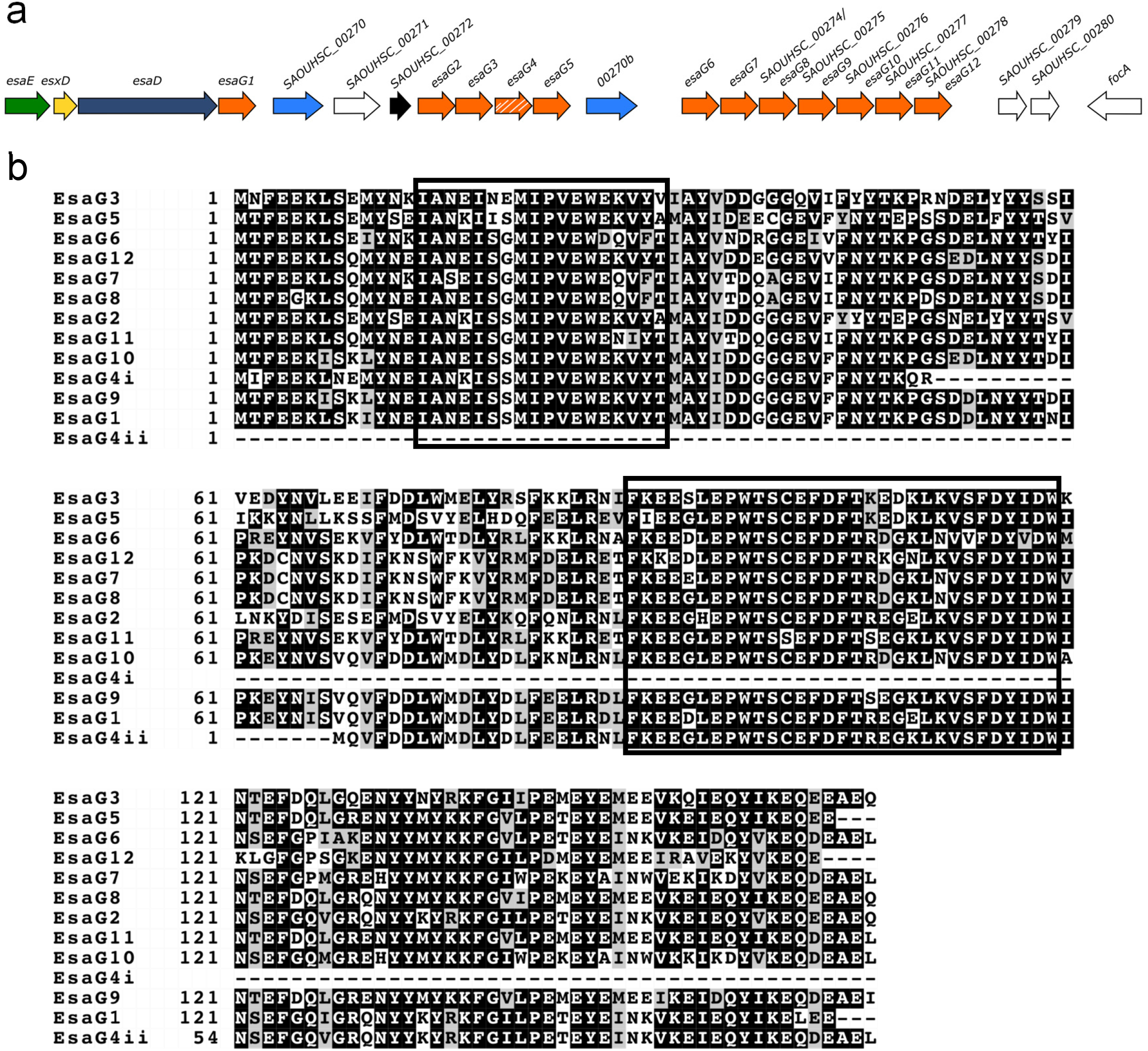
Homologues of *esaG* encoded at the *ess* locus in RN6390. a. Illustration of the 3’ of the RN6390 *ess* locus which encodes the T7SS nuclease toxin, EsaD, its cognate immunity protein, EsaG and 11 further homologues of EsaG (numbered *esaG2* – *esaG12*). Note that *esaG4* is shown in hatched shading because it is annotated as a pseudogene. However, it does encode two predicted ORFs, EsaG4i and EsaG4ii. b. Sequence alignment of EsaG homologues encoded by RN6390. The black boxes represent regions of high sequence similarity based on this alignment.

To assess variation between the 12 homologues of EsaG, the amino acid sequences were aligned (Fig 1b), and regions of high sequence similarity between the proteins were analysed using Plotcon (Fig 2a). Numerous regions of sequence similarity were observed, including two major blocks of high sequence homology at amino acids 13-31 and 84-119. To assess whether these were general features of *S. aureus* EsaG proteins, we downloaded approximately 4,000 *S. aureus* EsaG sequences from RefSeq and collectively analysed them using Plotcon. Fig S1a indicates that a similar profile of sequence similarity is seen across all EsaG proteins.

**Figure 2.**
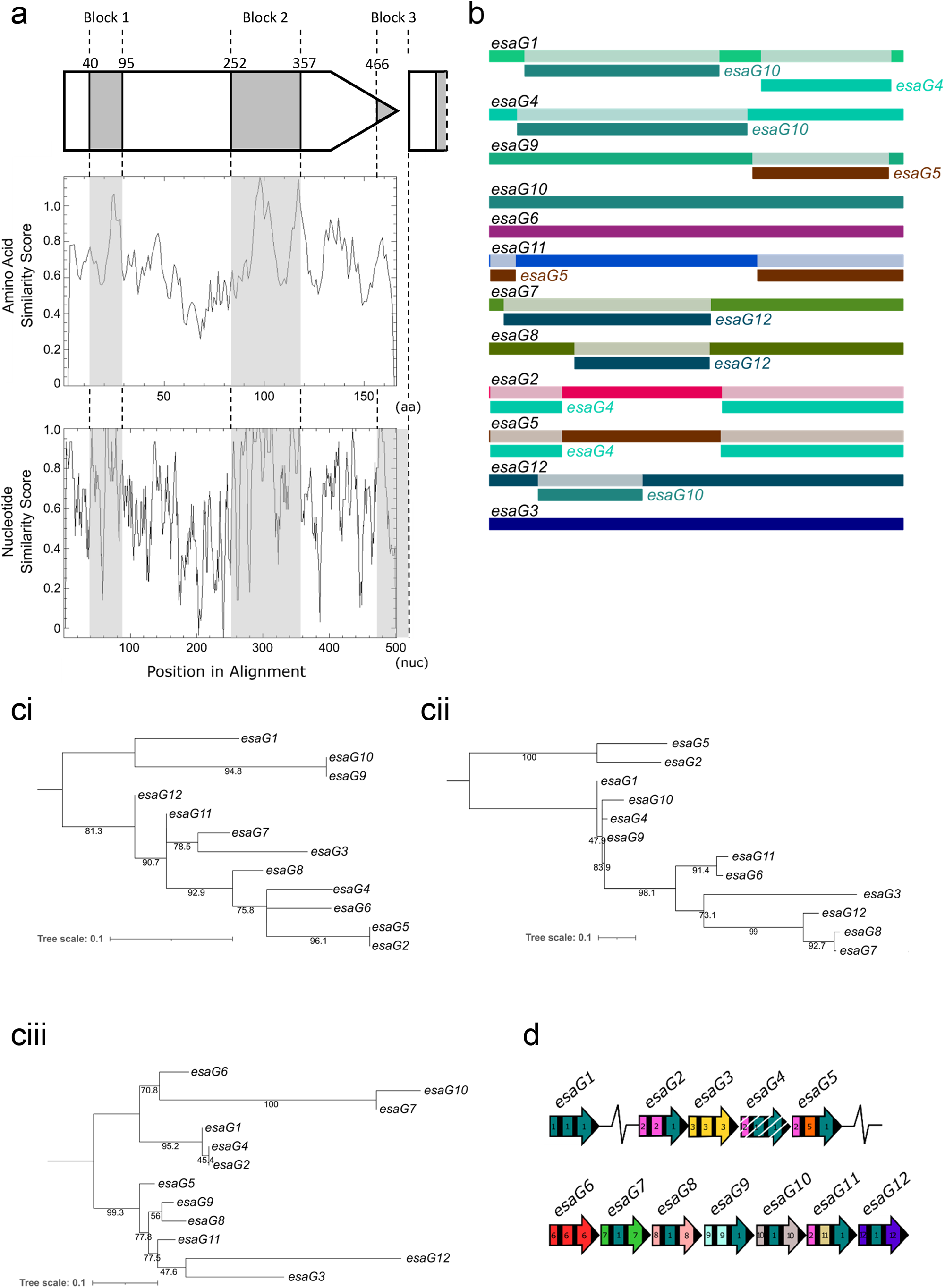
Recombination within the RN6390 *esaG* homologues. a. Regions of high sequence similarity across the RN6390 EsaG protein sequences (middle panel) and the corresponding nucleotide sequences (bottom panel). The positions of homology blocks are shown in grey shading and along with their relative positions along the gene sequence (top panel). The basepair positions that define the conserved regions are taken from the nucleotide sequences of *esaG1*. b. RDP4 was used to predict recombination events within the *esaG* homologues encoded by RN6390. The identity of each gene is given in black at the left, with regions of recombination labelled directly below, in the colour of the gene from which the recombinant section originated. ci-iii. For each of the three variable regions of the RN6390 *esaG* homologues a maximum likelihood tree was generated in IQTREE and visualised and annotated in iTOL. The variable regions consist of nucleotide 1-39, 96-251 and 358-465 of *esaG* respectively d. Illustration of regions of high sequence similarity in the *esaG* homologues in RN6390. Black bars represent conserved regions of the gene and the variable regions have been assigned a colour and corresponding number. Homologous regions are coloured with the same colour. Numbers were assigned based on the first gene in the series that had the unique variable region. White hatched shading indicates a pseudogene.

We next examined the intergenic regions between the RN6390 *esaG* homologues. Strikingly, we noted that most of them were of a very similar length, other than when they directly preceded a non-*esaG* gene (for example *SAOUHSC_00270b*). They also share a high degree of sequence similarity (Fig S2a). When we undertook Plotcon analysis on the *esaG* genes, including the 3’ intergenic regions (Fig 2a, Fig S2b), we noted the same two major homology blocks that we had seen from the amino acid analysis, but in addition a third block encompassing the end of the gene and the downstream intergenic region (Fig 2a, Fig S2b, Fig S2c).

Given the substantial levels of sequence similarity between the *esaG* genes, we used the recombination prediction software, RDP4, to determine whether there had been recombination events between the genes. The RDP4 output, shown in Table S1 and summarised Fig 2b, predicts with high significance that there have been extensive recombination events within most (but not all) of these genes. As shown in Fig 2b, recombination appears to occur at the three points within the genes that correspond to the regions of high nucleotide sequence homology. To support these findings, we constructed a maximum-likelihood tree for each of the variable regions of *esaG1-12* (Fig 2ci-iii). The *esaG* genes do not cluster in the same way in each variable region, confirming that these genes have not evolved solely by divergent evolution. For example, *esaG7* clusters with *esaG3* in variable region 1, *esaG8* in variable region 2 and *esaG10* in variable region 3 (Fig 2ciii). This suggests extensive recombination has occurred between these genes to generate this repertoire.

Based on the RDP4 results, we built a schematic representation showing the homologous regions of the *esaG* genes that are likely involved in the recombination events (Fig 2d). Specific regions of the genes seem to share high sequence similarity with others, for example, the mid-section of many of the homologues share high similarity to equivalent sections of *esaG1* (coloured teal). Conversely, genes such as *esaG3* and *esaG6* appear to be much more diverse.

### Evidence of *esaG* recombination in the USA300 epidemic strains of *S. aureus*

To determine whether recombination could be detected between *esaG* genes across *S. aureus* strains, we first examined genome sequences from a recent study analysing the community spread and evolution of a USA300 variant during a New York outbreak (32). USA300 is a methicillin-resistant *S. aureus essC1* strain, and a dominant cause of community-acquired *S. aureus* infection in the USA (33). Comparing the whole-genome sequences of the epidemic lineage showed that these strains carry only seven *esaG* genes in comparison to the ten copies in the closely related USA300 FPR3757. Using strain BKV_2 as a representative of the outbreak lineage, we aligned the region spanning from *esaE* to *SAUSA300_0303* with USA300 FPR3757. The alignment showed almost complete sequence identity, other than in a region spanning from the middle of *SAUSA300_0295* to the middle of *SAUSA300_0299* (Fig 3a) which was absent from BKV_2. When the nucleotide sequences of *SAUSA300_0295* and *SAUSA300_0299* were aligned with *esaG4* of BKV_2, *esaG4* was seen to be mosaic, with most of the gene being identical to *SAUSA300_0295* but the 3’ end showing 100% identity to *SAUSA300_0299* (Fig 3b). This is consistent with recombination between homology block 2 of *SAUSA300_0295* and *SAUSA300_0299*, with loss of the intervening DNA.

**Figure 3.**
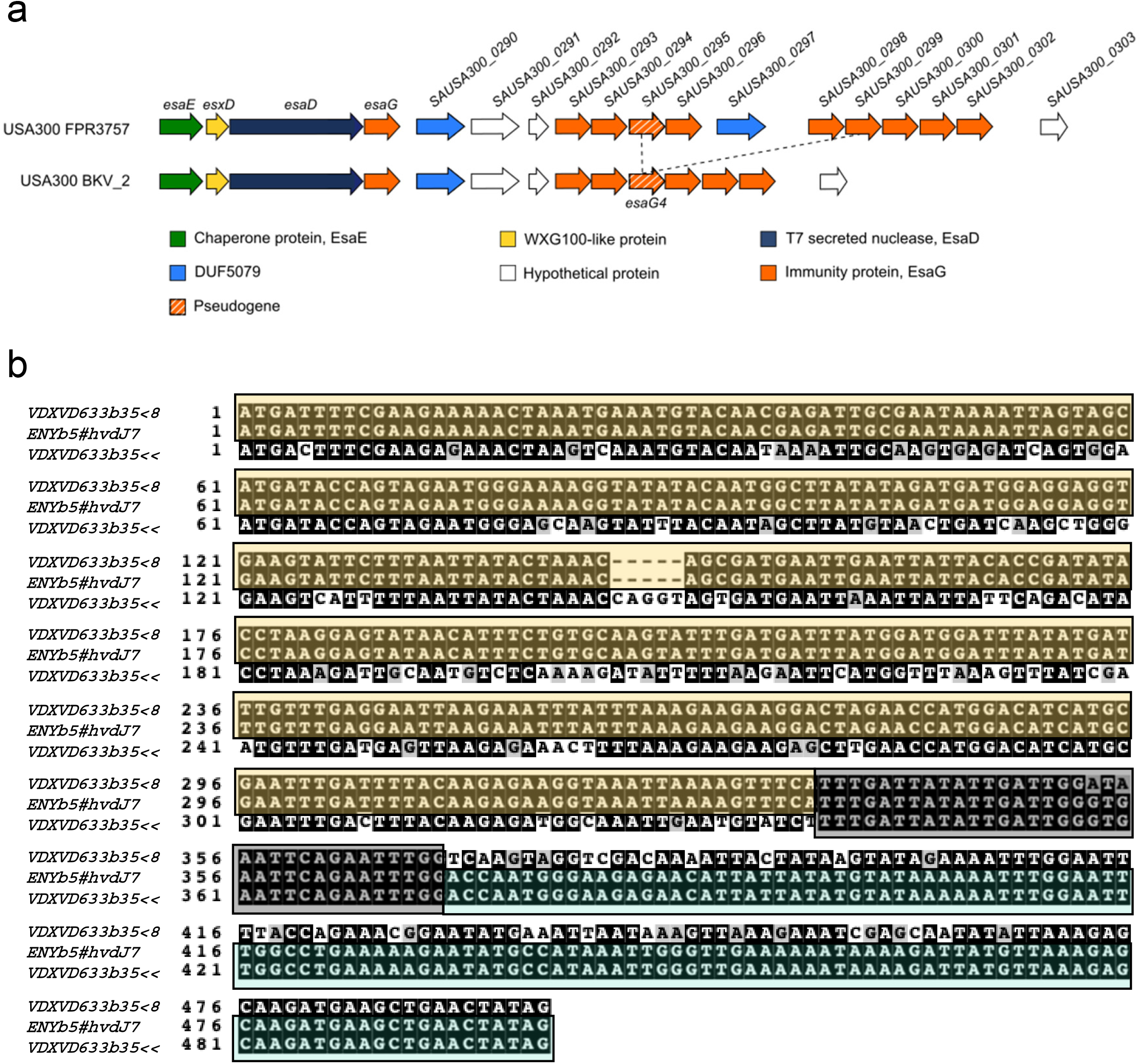
A recombination event in an epidemic lineage of USA300 results in loss of part of an *esaG* cluster and generation of a novel *esaG* gene. a. The *esaD* locus of USA300 FPR3757 and USA300 BKV_2. The dashed lines represent the region that is missing from the epidemic strain, USA300 BKV_2, when compared to the USA300 FPR3757 type strain. b. Nucleotide sequence alignment for *SAUSA300_0295* and *SAUSA300_0299* from USA300 FPR3757 and *esaG4* from USA300 BKV_2. Coloured blocks indicated homology between *BKV_2 esaG4* and the genes with which it is aligned.

### Analysis of *esaG* recombination within a historical culture collection

To obtain further evidence of *esaG* recombination events, we accessed genome sequences from the National Collection of Type Cultures (NCTC) which has recently generated long read assemblies from 133 *S. aureus* strains sampled over the past 100 years. From this resource we initially constructed a dataset of all the genome sequences (including NCTC8325 which is part of this collection). The coordinates of the *ess/T7* locus were identified in each strain using a custom bash script and the locus extracted using the coordinates for DNA immediately 5’ of *essC* and 3’ of *focA* (which defines the 3’ boundary of the T7 immunity island (10); Supplementary Data 1). Next the clonal complex of each strain was annotated based on a core genome SNP tree constructed previously (34), because for the purpose of our analysis it was essential to analyse very closely related strains to ensure that any differences could not be due to divergent evolution. From the annotated tree we selected to analyse the *ess/T7* locus from the 31 CC8 strains as there are a significant number of strains in this clonal complex (including NCTC8325) and all strains are extremely closely related to one another (34). Using Easyfig to assess the genetic differences at this locus (35), we observed that the most common genetic composition was to encode 12 *esaG* paralogues, as seen for NCTC8325 and RN6390 (Fig S3). However, there were clear differences in the *esaG* repertoire among some of these closely related strains, including two unique expansion events, five unique deletion events and two instances of recombination occurring in other regions of the locus (Fig S3). These were confirmed by long read sequence mapping (Supplementary Data 2).

Two unique expansion events were observed resulting in an additional *esaG7*-like gene in strain NCTC10724 and a duplication of *esaG11* in NCTC10702. Further analysis of the events in NCTC10724 indicate that recombination has occurred at homology block 1 of *esaG7* with homology block 1 of *esaG8*, most likely during DNA replication (Fig 4a). The duplication of *esaG11* in NCTC10702 is most likely due to recombination at homology block 3 between the 3’ of *esaG11* and the 3’ of *esaG10* (Fig 4b). These observations confirm that expansion of *esaG* repertoires can occur due to recombination between the homology blocks we have identified.

**Figure 4.**
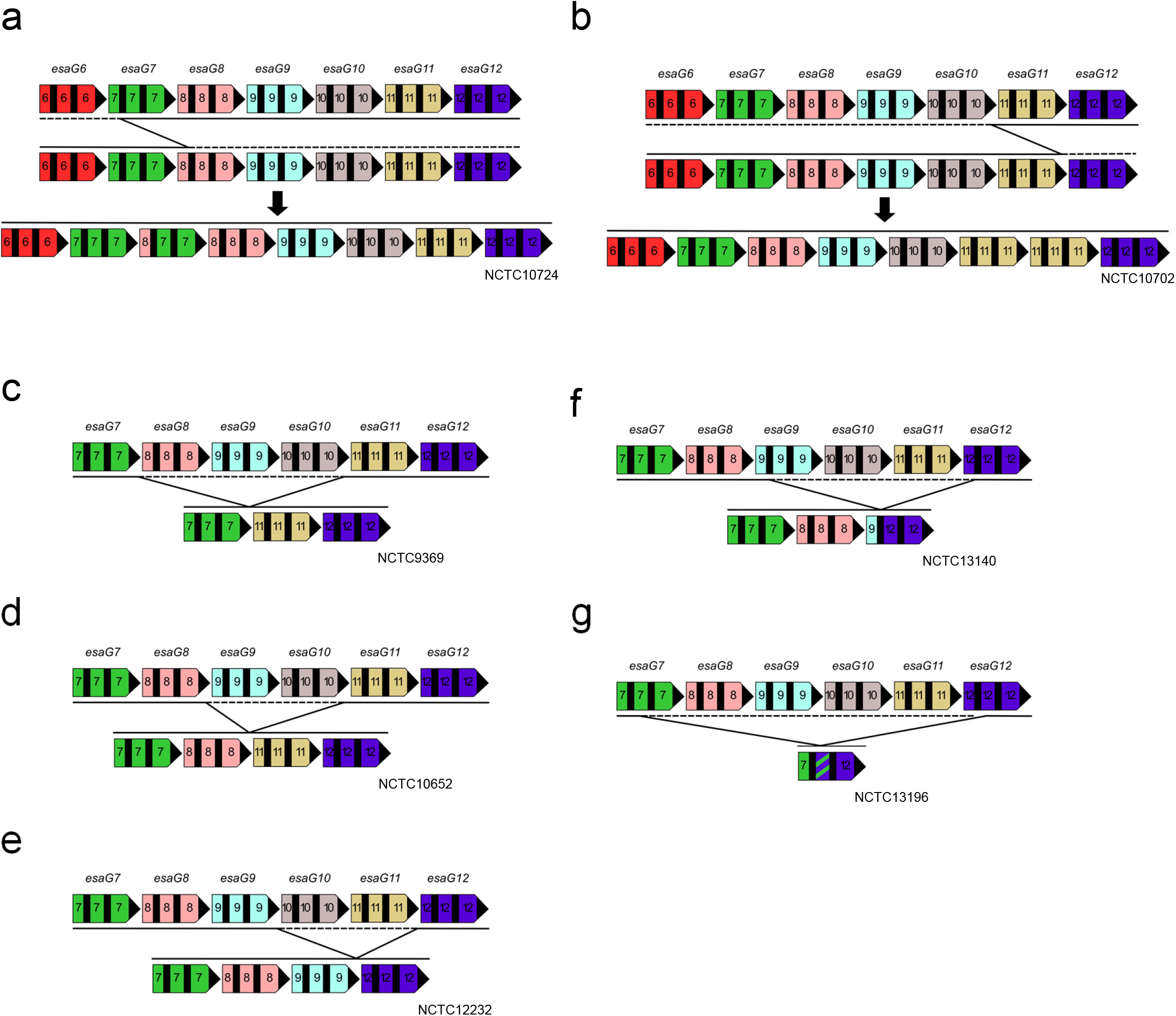
Recombination between *esaG* paralogs results in variation at the *ess* locus in CC8 *S. aureus* strains from a historical culture collection. Two expansion events of the *esaG* repertoire in the NCTC CC8 dataset were identified in a. NCTC10724 and b. NCTC10702 as respective representatives. The solid black line depicts the predicted recombination event to give the recombinant sequence depicted below. Five unique loss events were also identified in this dataset, represented by strains c. NCTC9369, d. NCTC10652, e. NCTC12232, f. NCTC13140 and g. NCTC13196. The dashed line represents the region deleted in the recombinant. Genes were given an individual colour and numbered based on *esaG1-12*, to make clear any recombination events.

Five unique deletion events were also identified among the strains. Deletions due to recombination at homology block 3 were responsible for three of these (Fig 4c-e). Recombination between homology block 1 of *esaG9* and *esaG12* resulted in a deletion event in NCTC13140, generating a novel *esaG* mosaic (Fig 4f). In NCTC13196, a deletion event has occurred to produce a novel *esaG* mosaic comprising the 5’ part of *esaG7* and 3’ end of *esaG12*. Notably in this instance, recombination in these genes has not occurred within the homology blocks, but instead, in variable region 2 of the gene (Fig 4g). This is possible because *esaG7* and *esaG12* share high sequence similarity in this region (Fig S2c, Fig 2d).

In RN6390 and other CC8 strains carrying the full repertoire of 12 *esaG* homologs, two *DU5079* genes are present, one encoded immediately downstream of *esaG1* and the other downstream of *esaG5* – these genes are annotated *DUF5079-1 (SAOUHSC_00270)* and *DUF5079-2 (SAOUHSC_00270b)*, respectively (Fig 5). In ten of the CC8 strains analysed, we noted that only a single *DUF5079* gene was encoded, and that these strains carried only *esaG1* along with *esaG6* – *esaG12*. Analysis reveals that rather than a recombination event between homology block 3 of *esaG1* and *esaG5*, recombination was instead highly likely to have occurred between the *DU5079* regions. In strain NCTC10702, recombination occurred between *DUF5079-1* and *DUF5079-2* to produce a recombinant *DUF5079* gene (Fig 5a), whereas in strain NCTC12232 only the *DUF5079-2* gene was found, suggesting recombination had occurred between the intergenic regions upstream of these genes (Fig 5b). Aligning the region spanning *esaG1* and *DUF5079-1* with *esaG5* and *DUF5079-2* from RN6390, we noted that the intergenic region and *DUF5079* gene share much higher sequence similarity than most of the *esaG* genes (Fig 5c). To explore this further, these genomic regions were extracted from the CC8 collection and an alignment visualised using Plotcon, confirming that across these strains, high sequence similarity is observed within the intergenic regions and *DUF5079* genes (Fig 5d). This indicates that recombination in other regions of the *ess/T7* locus also plays a role in dictating the repertoire of T7SS immunity genes carried at this locus.

**Figure 5.**
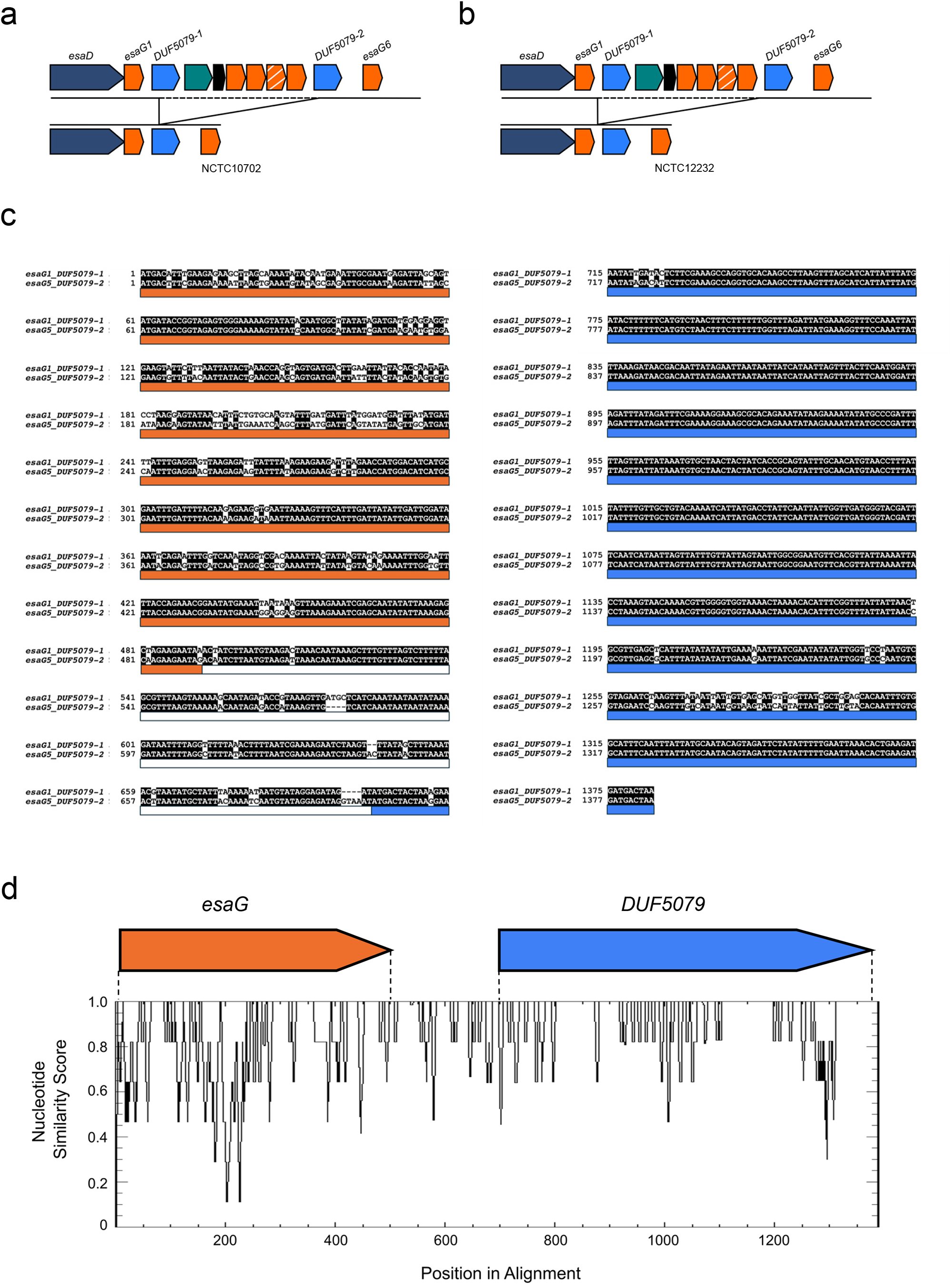
Recombination also occurs between *DUF5079* genes. Recombination events were detected between the two *DUF5079* genes at the *ess* locus in the NCTC *S. aureus* CC8 dataset. Recombination occurred a. within the *DUF5079* genes and b. in the intergenic regions directly upstream of these, resulting in the loss of *esaG2-5* in several strains. c. An alignment of the region spanning the 5’ of *esaG1* to the 3’ of *DUF5079-1* with the 5’ of *esaG5* to the 3’ of *DUF5079-2*. Coloured bars below the alignment represent *esaG* (orange), the intergenic region (white) and *DUF5079* (blue). d. All instances of the region spanning the 5’ of *esaG* to the 3’ of *DUF5079* was manually extracted from the NCTC CC8 dataset, aligned using MAFFT and visualised using Plotcon to identify regions of high similarity.

### *esaG* diversity across *S. aureus essC2, essC3* and *essC4* variants

Although *S. aureus essC2, essC3* and *essC4* variants do not encode EsaD, these strains all accumulate *esaG* genes at the 3’ end of their *ess/T7* loci. To ascertain whether homologous recombination may similarly play a role in the evolution of these *esaG* families we analysed the *esaG* genes encoded in strains ST398, MRSA252 and HO 5096 0412, as representatives of *essC2, essC3* and *essC4* variants, respectively (Fig S4). We noted that none of the *esaG* genes found in these strains appear to be direct copies of the 12 *esaG* genes we identified in the CC8 *essC1* strains (Fig S4 a-d). Using RDP4 analysis to detect recombination events, again we could detect clear signatures of recombination in almost all of the genes (Fig S4 e-g; Table S2-4), with the exception of *SAPIG_0314-0315* pseudogene. We conclude that recombination events are common among *esaG* genes in all *S. aureus* strains analysed.

### No clear evidence for recombination within *tsaI* genes

TspA is a T7SS-secreted antibacterial toxin that is highly conserved across all *essC* variant strains (19). In all strains it is encoded away from the T7SS gene cluster, at a genomic location bounded by *SAOUHSC_00583* and *iolS* (*SAOUHSC_00603*). The toxic activity of TspA is neutralised by TsaI, a membrane protein of the DUF443 family (19). Multiple copies of *tsaI* genes are encoded downstream of *tspA* (Fig S5a,b), and in RN6390 there are 11 copies, *tsaI1* – *tsaI11* (Fig S5a). A small pseudogene, encoded by *SAOUHSC_00600* shares high sequence similarity to part of the toxin region of TspA and is also found at this locus.

To determine whether recombination events also contributed to the evolution of the TsaI repertoires, we aligned the amino acid sequences of RN6390 TsaI proteins (Fig S5c). We noted much greater sequence variability between these proteins than the EsaG homologues, particularly in the C-terminal region of the protein. Similar variability was observed in a representative alignment of around 3000 TsaI sequences (Fig S1b). At the nucleotide sequence level, much greater variability was also observed in the *tsaI* intergenic regions, in both length and DNA sequence compared with *esaG* (Fig S5d). Using Plotcon we identified only a single region of high similarity among TsaI sequences, covering approximately the first 75 amino acids, which is also mirrored at the DNA level (Fig S6a). As only one block of high sequence similarity is detected and there is a high degree of sequence variability in the *tsaI* intergenic regions, recombination within individual genes is unlikely. To analyse this, we used RDP4 to predict recombination events within the 11 *tsaI* genes (Fig S6b). Far fewer potential recombination events were predicted than for *esaG* genes, and with much lower probability (Table S5). We conclude that evolutionary processes other than tandem homologous recombination give rise to the variable *tsaI* gene clusters in *S. aureus* genomes.

### Intergenic recombination between *tipC* immunity genes in *Streptococcus*

Three T7SS-secreted antibacterial toxins have been identified in *Streptococcus intermedius*. TelA and TelB are both cytoplasmic-acting toxins neutralised by the TipA and TipB immunity proteins, respectively (12). Our genome analysis indicates that strains generally encode only a single *tipA* gene, while *tipB* is found in up to five copies. The third *S. intermedius* toxin is TelC, a lipid II phosphatase. Protection from TelC toxicity is provided by TipC, a membrane-bound immunity protein that faces the extracellular space (12, 36). Klein *et al*. reported that the number of *tipC* genes encoded at the *telC* locus was highly variable between strains (36). We used gene neighbourhood analysis across Streptococcal genomes to compare the number of *tipC* genes present at the *telC* gene cluster, identifying 15 of them in a strain of *Streptococcus mitis* BCC08 (Fig 6a).

**Figure 6.**
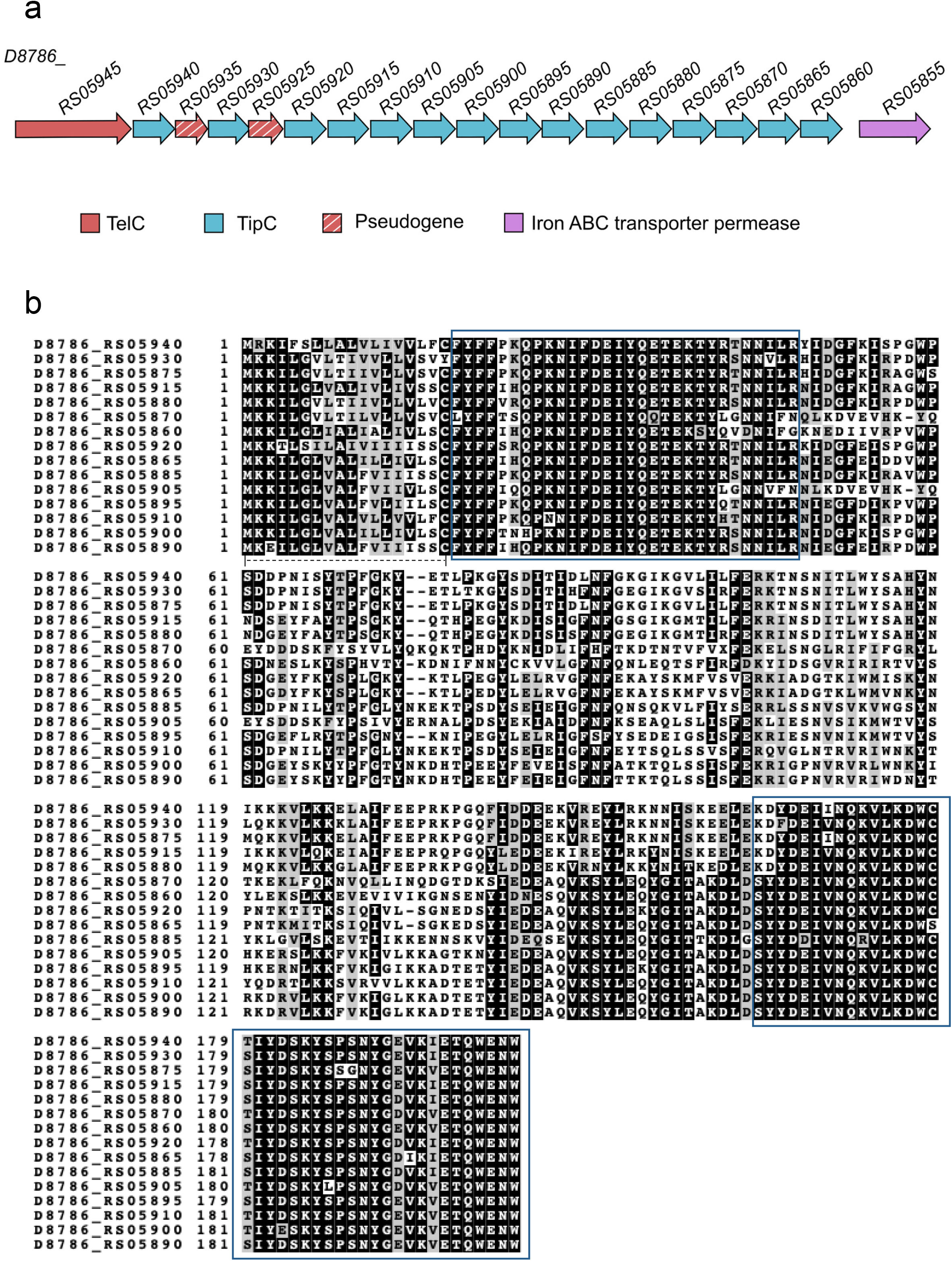
Homologues of *tipC* encoded at the *telC* locus of *Streptococcus mitis* BCC08. a. Genetic arrangement of *tipC* genes in *S. mitis* BCC08. b. An alignment of the encoded TipC homologues. The blue boxes represent regions of high sequence similarity, and the dashed line at the N-terminus of the aligned sequences indicates a probable lipoprotein signal peptide.

Alignment of the *S. mitis* BCC08 TipC sequences and their encoding DNA (Fig 6b, Fig S7) showed two regions of high sequence conservation close to the start and end, with a central region of much higher sequence variability (Fig 7a). Using RDP4 to screen for recombination, at least five recombination events were predicted between these genes (Fig 7b, Table S6), in each case almost certainly through the homology blocks we identified. To analyse this further we constructed a maximum likelihood tree to compare *tipC* gene sequences. Genes *D8786*_*RS05910* and *D8786*_*RS0585* cluster closely in this tree, which corresponds to the recombination event predicted between these two genes (Fig 7b, Table S6). Likewise, *D8786*_*RS05920*, which is predicted to be the major parent to *D8786*_*RS05865* (Table S6) also clusters phylogenetically with this gene. We conclude that similar to *esaG*, intergenic recombination drives the evolution of *tipC* immunity gene repertoires.

**Figure 7.**
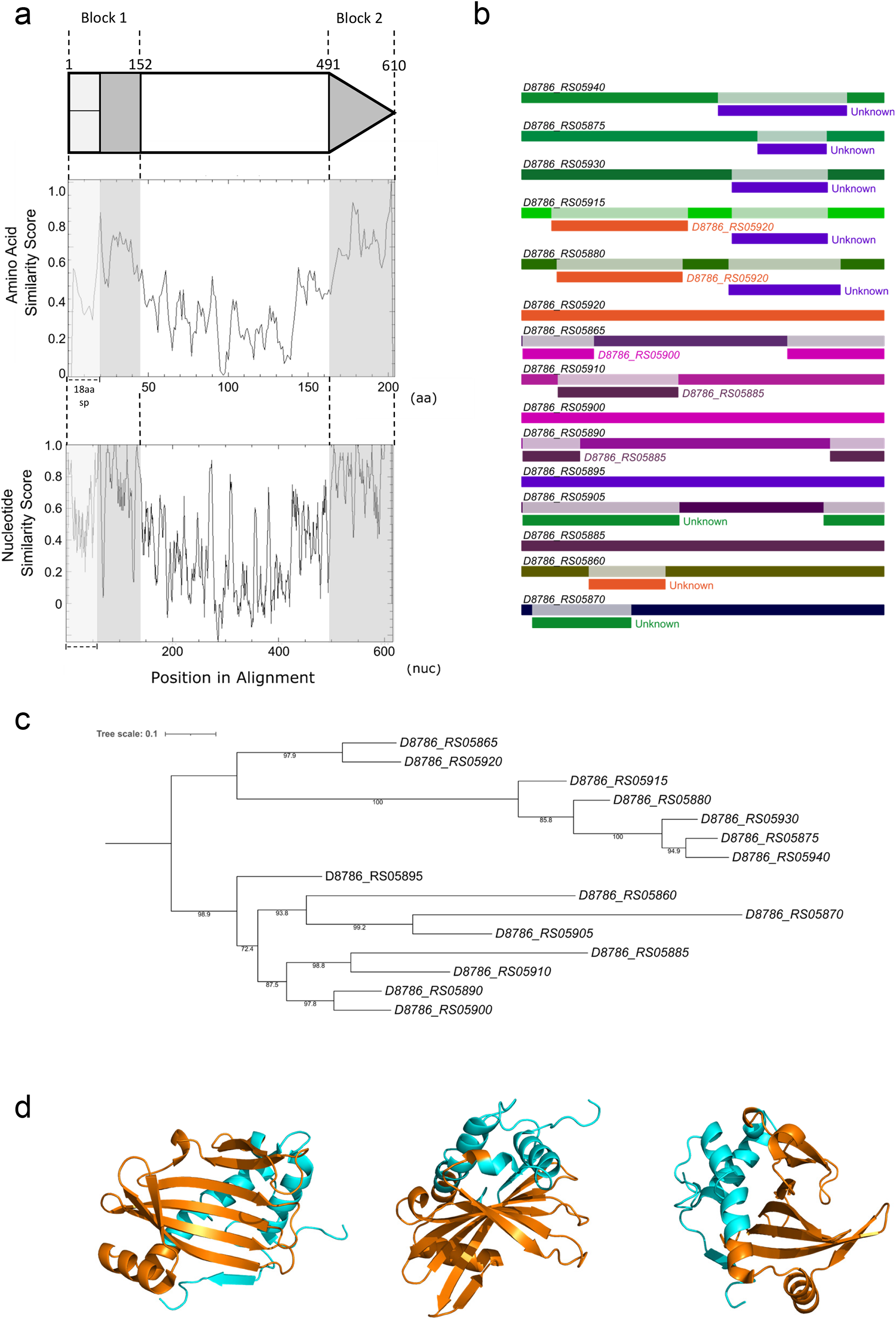
Recombination within the *S. mitis* BCC08 *tipC* homologues. a. Regions of high sequence similarity across the *S. mitis* BCC08 TipC protein sequences (middle panel) and the corresponding nucleotide sequences (bottom panel). The positions of homology blocks are shown in grey shading, with their relative positions along the gene sequence (top). The first 18 amino acids of TipC form a predicted lipoprotein signal sequence which is indicated by pale grey shading. The basepair positions that define the conserved regions are taken from the nucleotide sequences of *D8786_RS05940*. b. RDP4 was used to predict recombination events within the *tipC* homologues. The identity of each gene is given in black at the left, with regions of recombination labelled directly below, in the colour of the gene from which the recombinant section originated. c. A maximum likelihood tree was generated for *tipC* homologues in IQTREE and visualised and annotated in iTOL. d. The conserved (cyan) and variable (orange) regions of TipC were mapped to the crystal structure of *S. intermedius* TipC2 (pdb:6DHX; 53).

## DISCUSSION

In this study we have undertaken sequence analysis of *S. aureus* EsaG proteins and their encoding DNA, including their 3’ flanking regions, to investigate how copy number variability may arise. We found three blocks of highly conserved nucleotide sequence, a large central one of approximately 100 nucleotides in length, and a 5’ and 3’ block both of around 55 nucleotides each. Homologous recombination occurs at regions of high similarity within nucleotide sequences (Reviewed in 37). The minimum length requirement for efficient recombination in *S. aureus* is unclear but stretches of 40-70 nucleotides have been reported for other bacteria (38-40). Using RDP4 to predict recombination within *esaG* genes, we found strong evidence for recombination, corresponding to events within the three homology blocks we identified. Through the analysis of genome sequences from multiple closely related CC8 strains we found extensive evidence of recombination within *esaG* genes to alter copy number, and in some instances the sequence of individual *esaG* genes.

Previous work has reported that the *S. aureus* tandem-like lipoproteins, encoded on the νSaα island, also show extensive copy number variation across strains (41, 42). Similar analysis to that reported here showed that each *lpl* gene shares a stretch of approximately 130 nucleotides of high sequence similarity in its central region. Recombination was demonstrated to occur between the central conserved region of one gene and the same region of the neighbouring gene (40), thus spanning the 3’ portion of gene 1, the intergenic region and the 5’ region of gene 2.

To investigate whether intergenic recombination might represent a general mechanism for the evolution of T7SS immunity gene families, we examined the organisation of the *tipA, tipB* and *tipC* immunity genes in Streptococci. It has previously been noted that *tipC* copy number is highly variable in Streptococcal genomes (36), and our analysis identified that up to 15 copies of *tipC* could be present. Examination of recombination events within *tipC* revealed that intergenic recombination is also a feature, and that it primarily occurs between homologous blocks of sequence identity at the start and end of the genes. The structure of TipC reveals that it has seven beta strands forming a concave face, with three alpha helices made up from the N- and C-terminal regions of the protein (53, Fig 7d). Analysis of colicin DNase toxins and their immunity proteins has shown that sequence divergence between related toxins and immunities tends to concentrate at the binding interface (43). In agreement with this, site-directed mutagenesis has strongly implicated the concave face as the region that binds to TelC, and this region of TipC shows the highest level of sequence divergence (Fig 7d; 36). We speculate that for EsaG the regions of high sequence similarity are not directly involved in toxin binding but may provide a structural framework on which amino acid substitutions in the variable regions accumulate to alter the toxin binding specificity. Mechanisms to allow rapid evolution of immunity proteins are likely to be essential to allow strains to quickly acquire resistance to novel toxin variants. We have also demonstrated that strains can lose significant numbers of immunity genes from such repertoires. This would be expected to reduce the metabolic burden of synthesising a large number of similar proteins in an environment where they provide no benefit. It is likely that there is a trade-off between carrying sufficient immunity proteins to protect against competitors and the metabolic consequences of producing multiple protein variants.

TspA is a second polymorphic toxin encoded by *S. aureus*. Protection from its membrane-depolarising activity is provided by the immunity protein TsaI (19). TsaI is a polytopic membrane protein predicted to have five transmembrane domains. As with *esaG*, multiple *tsaI* genes are found in *S. aureus* genomes, however, our analysis has indicated that there is little evidence of recombination between them. We found only a single block of high nucleotide sequence similarity at the 5’ end of *tsaI* genes. This encodes approximately the first 75 amino acids of TsaI, which would encompass the first two transmembrane domains. At present it is not known whether TsaI neutralises TspA though direct interaction, or through the sequestering of a membrane-bound partner protein with which TspA must interact to facilitate its insertion or folding (or a combination of both). In this context, membrane permeabilising peptide bacteriocins produced by some Gram-positive bacteria require a membrane-bound receptor, such as the membrane components of the mannose phosphotransferase system, for their activity. In the case of the *Lactococcus lactis* lactococcin A bacteriocin, the immunity protein LciA acts through formation of a complex with both the receptor protein and the bacteriocin (44). By analogy it is possible that the first two transmembrane domains of TsaI interact with a candidate receptor, constraining their sequence, whereas the remainder of the immunity protein binds to the toxin and is therefore under diversifying selection. At present it is unclear how *tsaI* gene clusters evolve, although we did note that there was evidence for carriage of *tsaI* genes on Staphylococcal plasmids, which may facilitate gene movement within and between strains (data not shown). Further work would be required to clarify the mechanisms that drive *tsaI* diversity.

Fragments of a second copy of *tspA* are found within the *tsaI* gene cluster at the *tspA* locus in many (but not all) strains of *S. aureus*. Interestingly, an Rhs toxin encoded by *Salmonella enterica* serovar Typhimurium LT2 is also found as a second partial copy downstream of primary toxin locus; in each case both *rhs* copies precede an immunity gene. Repeated passage of the strain resulted in a fraction of the cells undergoing recombination between the *rhs* copies, generating a mosaic gene between *rhs1* and *rhs2* (45). An alignment of the *tspA* fragment (spanning the entire intergenic region between *tsaI10* and *tsaI11*) with full length *tspA* indicated that the regions of homology fall at the 3’ of *tspA*, although we did not find any evidence of recombination between these nucleotide sequences in the datasets we analysed. It should be noted that recombination between *tspA* and the *tspA* fragment would generate a novel sequence variant for this toxin, as well as deleting a cluster of *tsaI* genes lying between these regions, potentially helping to increase strain competitiveness under specific growth conditions.

## FUNDING INFORMATION

This work was supported by Wellcome through Investigator Award 110183/A/15/Z to T.P. and Sir Henry Wellcome Postdoctoral Fellowship 218622/Z/19/Z to G.M. The NCTC3000 project, which provided the 133 NCTC genome assemblies, was funded via Wellcome Trust grant 101503/Z/13/Z. S.R.G. is funded by the Newcastle-Liverpool-Durham BBSRC DTP2 Training Grant, project reference number BB/M011186/1.

### ACKNOWLEDGEMENTS

We thank Prof Jan-Maarten van Dijl (University of Groningen, NL) for providing us with strain RN6390, and Prof Victor Torres and Dr. Alejandro Pironti for providing the genome sequence of USA300 BKV_2.

## AUTHOR CONTRIBUTIONS

S.R.G. and T.P. conceptualised the study. S.R.G. carried out initial analysis. G.M. wrote custom scripts and produced these outputs. J.D. wrote custom scripts for the analysis of the NCTC dataset and provide the raw data for this. S.R.G. analysed the NCTC dataset. S.R.G. created the figures. S.R.G. and T.P. wrote the manuscript. All authors approved the final version of the manuscript.

