## Supplemental Figures for "Homologous recombination between tandem paralogues drives evolution of a subset of Type VII secretion system immunity genes in firmicute bacteria"

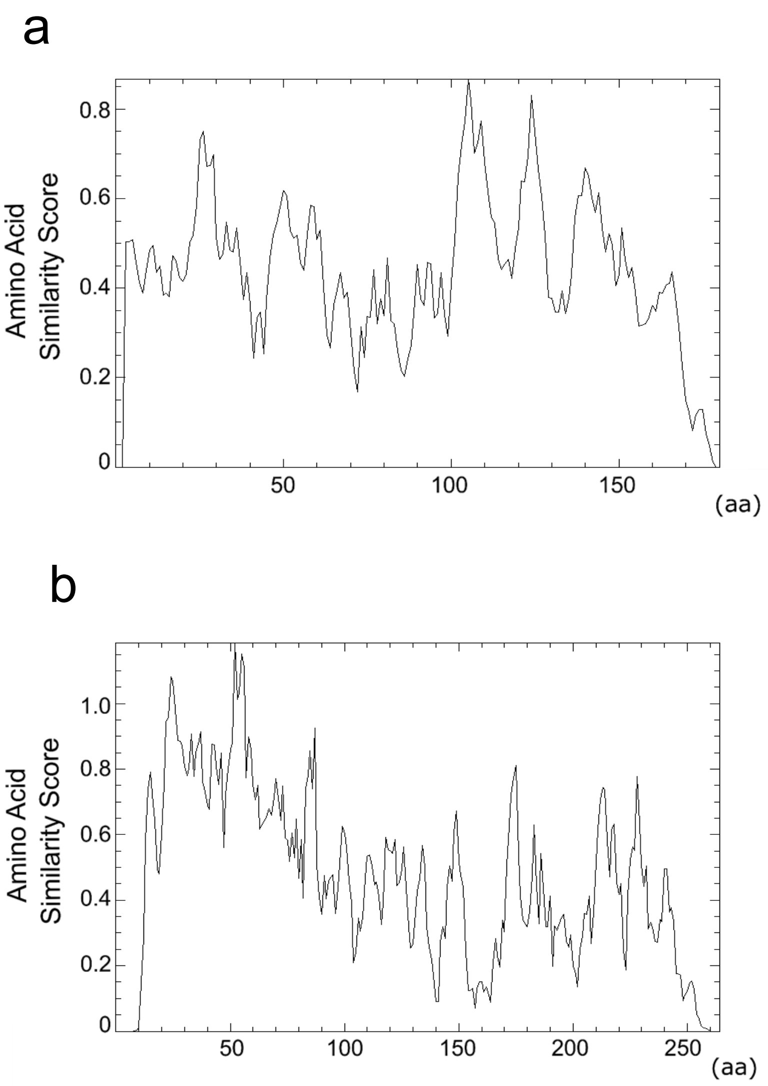


**Fig S1. Similarity plots for representative EsaG and TsaI amino acid sequences.** All available amino acid sequences for EsaG and TsaI were obtained from RefSeq. Sequences were aligned and a similarity plot produced using plotcon for a. EsaG and b. TsaI.

a

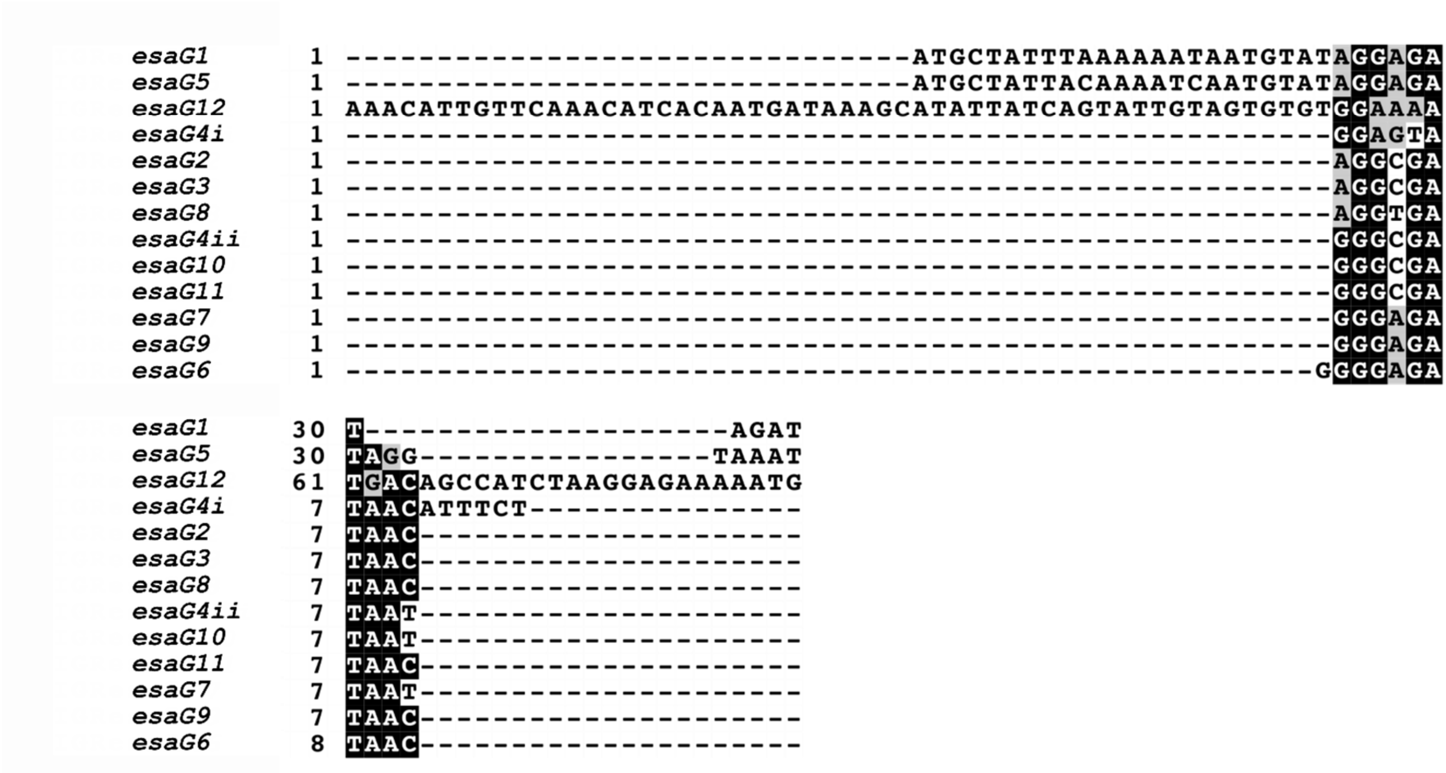

esaG1 1 -------------------------------ATGCTATTTAAAAAATAATGTATAGGAGA
esaG5 1 -------------------------------ATGCTATTACAAAATCAATGTATAGGAGA
esaG12 1 AAACATTGTTCAAACATCACAATGATAAAGCATATTATCAGTATTGTAGTGTGTGGAAAA
esaG4i 1 ------------------------------------------------------GGAGTA
esaG2 1 ------------------------------------------------------AGGCGA
esaG3 1 ------------------------------------------------------AGGCGA
esaG8 1 ------------------------------------------------------AGGTGA
esaG4ii 1 ------------------------------------------------------GGGCGA
esaG10 1 ------------------------------------------------------GGGCGA
esaG11 1 ------------------------------------------------------GGGCGA
esaG7 1 ------------------------------------------------------GGGAGA
esaG9 1 ------------------------------------------------------GGGAGA
esaG6 1 -----------------------------------------------------GGGGAGA
esaG1 30 T--------------------AGAT
esaG5 30 TAGG----------------TAAAT
esaG12 61 TGACAGCCATCTAAGGAGAAAAATG
esaG4i 7 TAACATTTCT---------------
esaG2 7 TAAC---------------------
esaG3 7 TAAC---------------------
esaG8 7 TAAC---------------------
esaG4ii 7 TAAT---------------------
esaG10 7 TAAT---------------------
esaG11 7 TAAC---------------------
esaG7 7 TAAT---------------------
esaG9 7 TAAC---------------------
esaG6 8 TAAC---------------------


b

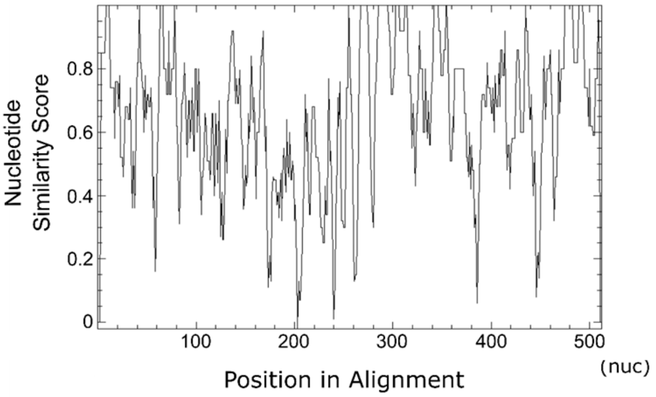

Nucleotide
Similarity Score
0
0.2
0.4
0.6
0.8
100
200
300
400
500
(nuc)
Position in Alignment

C

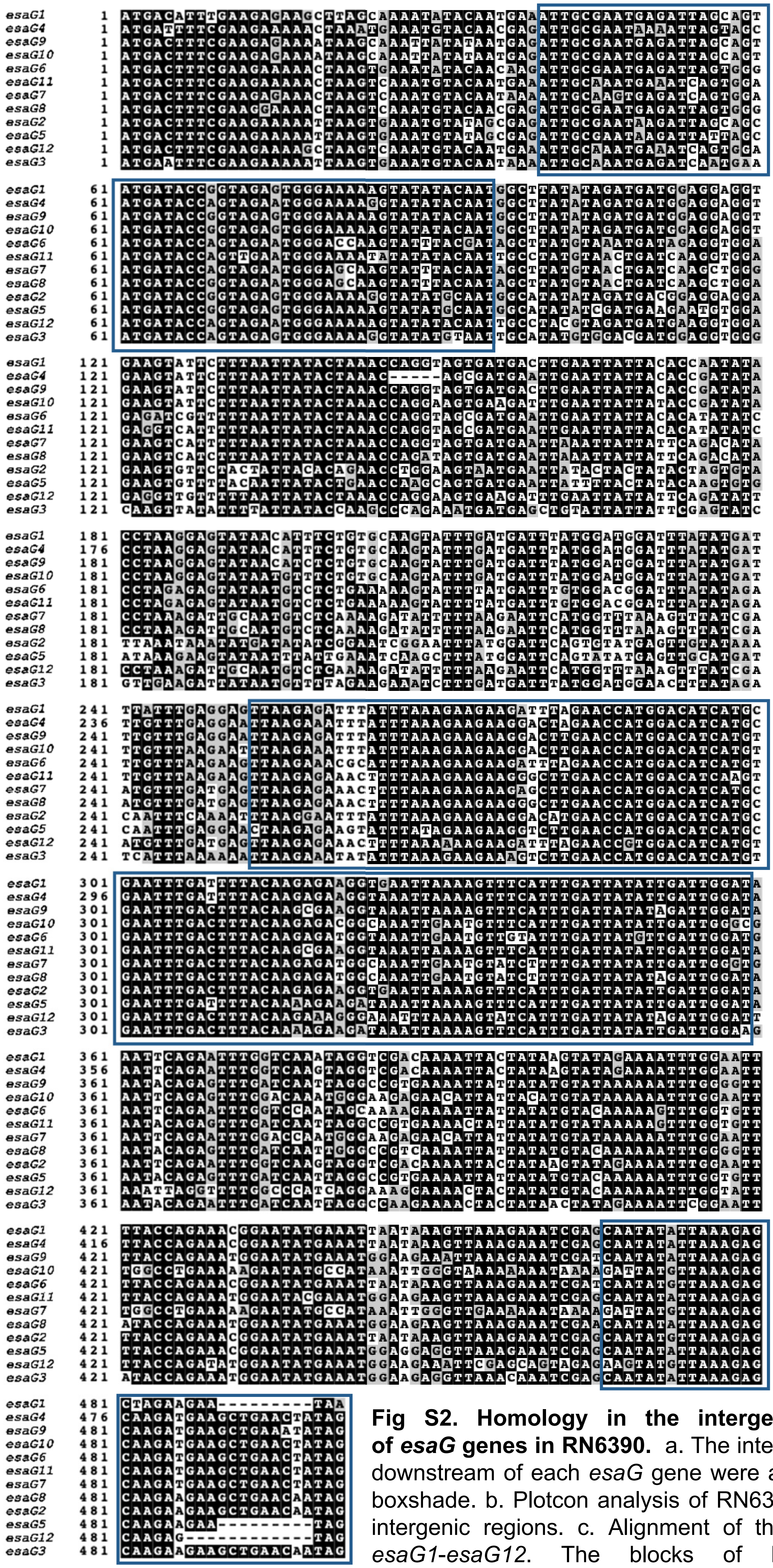

**Fig S2. Homology in the intergenic regions downstream of *esaG* genes in RN6390.** a. The intergenic regions found directly downstream of each *esaG* gene were aligned and visualised using boxshade. b. Plotcon analysis of RN6390 *esaG* genes and their 3' intergenic regions. c. Alignment of the nucleotide sequences of *esaG1*-*esaG12*. The blocks of high sequence similarity corresponding to Fig 2a are outlined in blue.

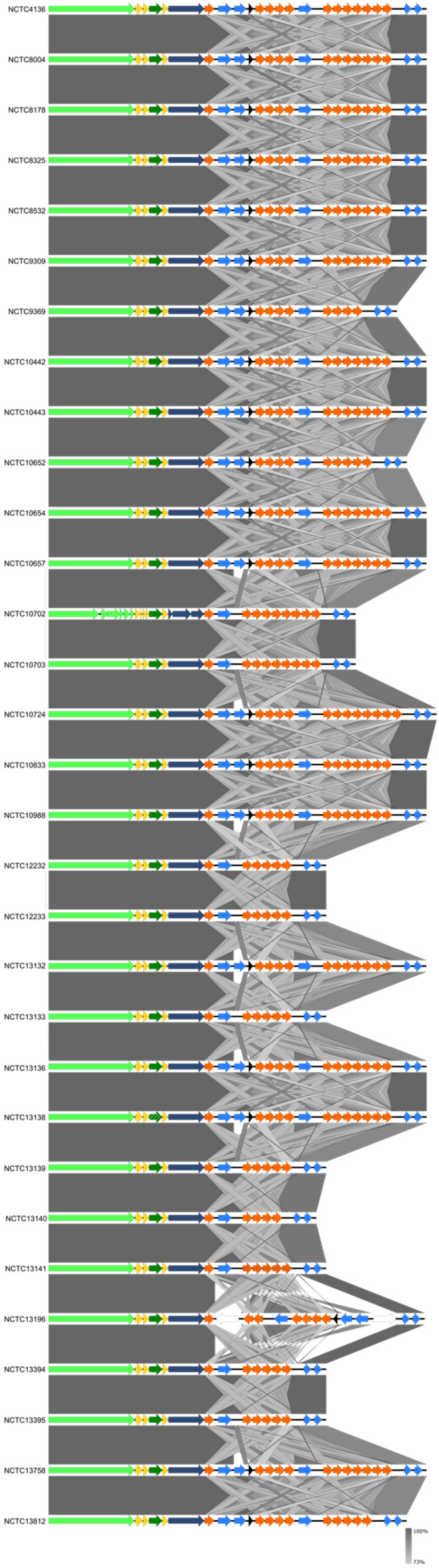


**Fig S3. Easyfig alignment of the *ess/T7* locus of CC8 strains from the NCTC culture collection.** The *ess/T7* locus was extracted from *S. aureus* strains from CC8 in the NCTC culture collection. Easyfig was used to perform pairwise alignment of strains, based on the order of accession in the NCTC collection.

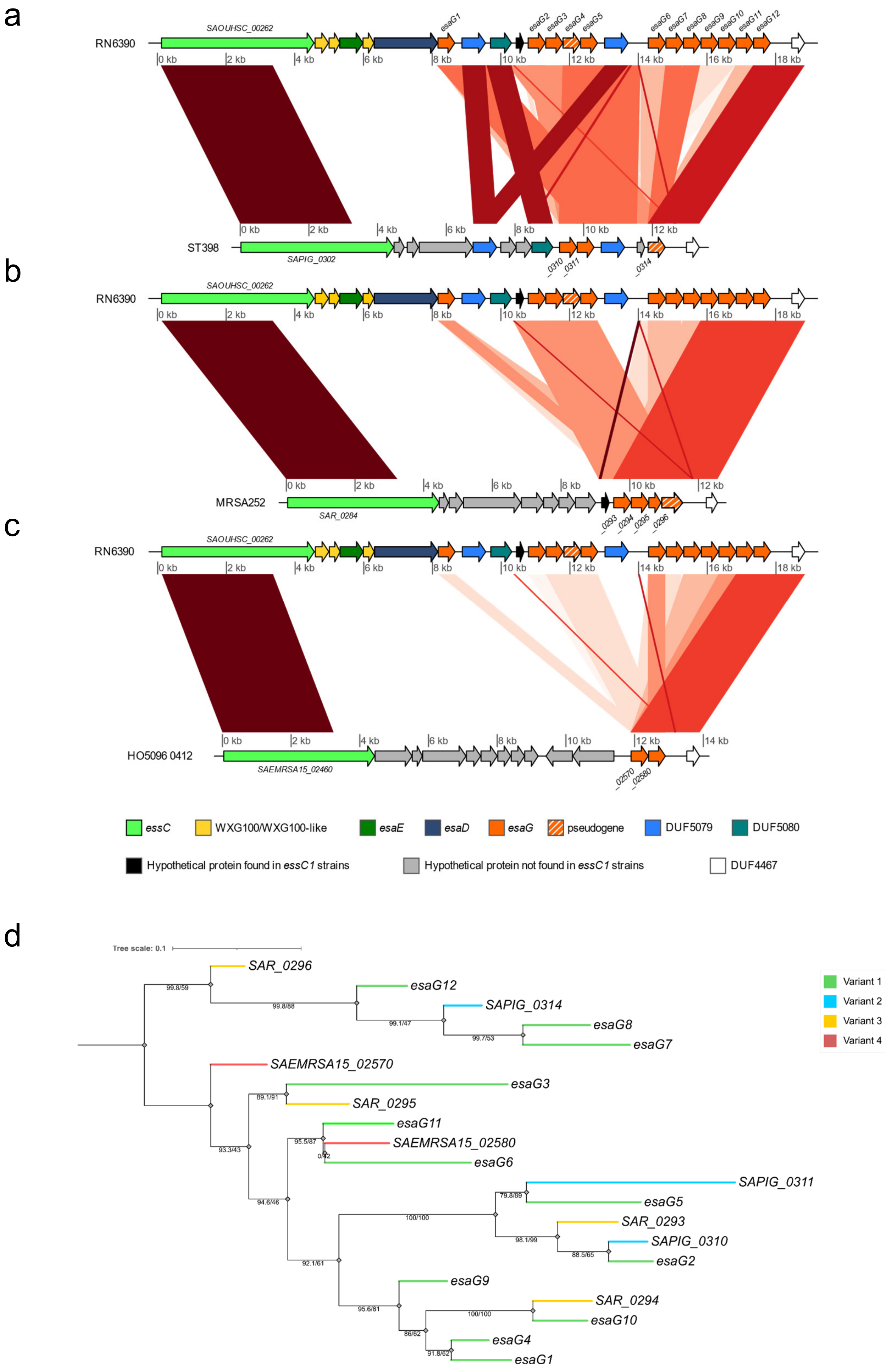
a
RN6390
SAOUHSC_00262
esaG1
esaG2
esaG3
esaG4
esaG5
esaG6
esaG7
esaG8
esaG9
esaG10
esaG11
esaG12
0 kb
2 kb
4 kb
6 kb
8 kb
10 kb
12 kb
14 kb
16 kb
18 kb
ST398
SAPIG_0302
_0310
_0311
_0314
b
MRSA252
SAR_0284
_0293
_0294
_0295
_0296
c
HO5096 0412
SAEMRSA15_02460
_02570
_02580
essC
WXG100/WXG100-like
esaE
esaD
esaG
pseudogene
DUF5079
DUF5080
Hypothetical protein found in essC1 strains
Hypothetical protein not found in essC1 strains
DUF4467
d
Tree scale: 0.1
SAR_0296
esaG12
SAPIG_0314
esaG8
esaG7
SAEMRSA15_02570
esaG3
SAR_0295
esaG11
SAEMRSA15_02580
esaG6
SAPIG_0311
esaG5
SAR_0293
SAPIG_0310
esaG2
esaG9
SAR_0294
esaG10
esaG4
esaG1
99.8/59
99.8/88
99.1/47
99.7/53
89.1/91
93.3/43
95.5/87
0/42
94.6/46
79.8/89
100/100
98.1/99
88.5/65
92.1/61
95.6/81
100/100
86/62
91.8/62
Variant 1
Variant 2
Variant 3
Variant 4

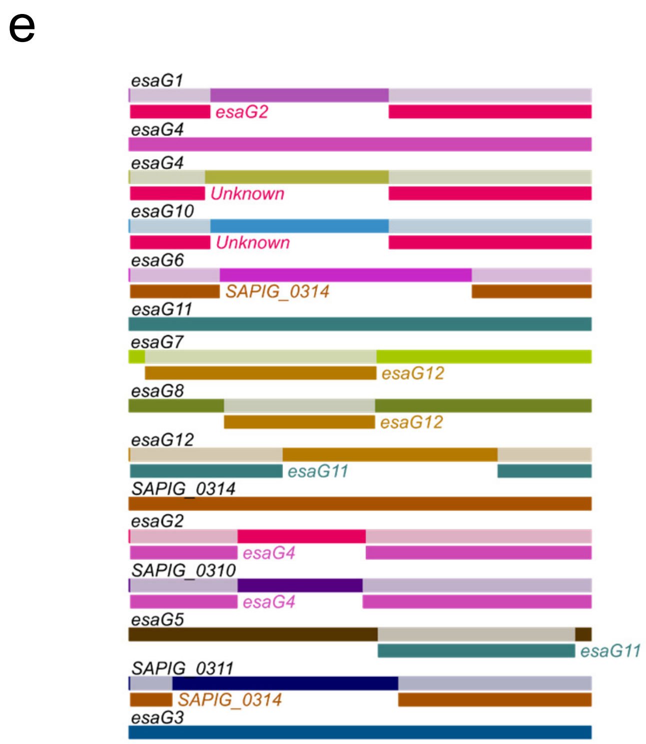


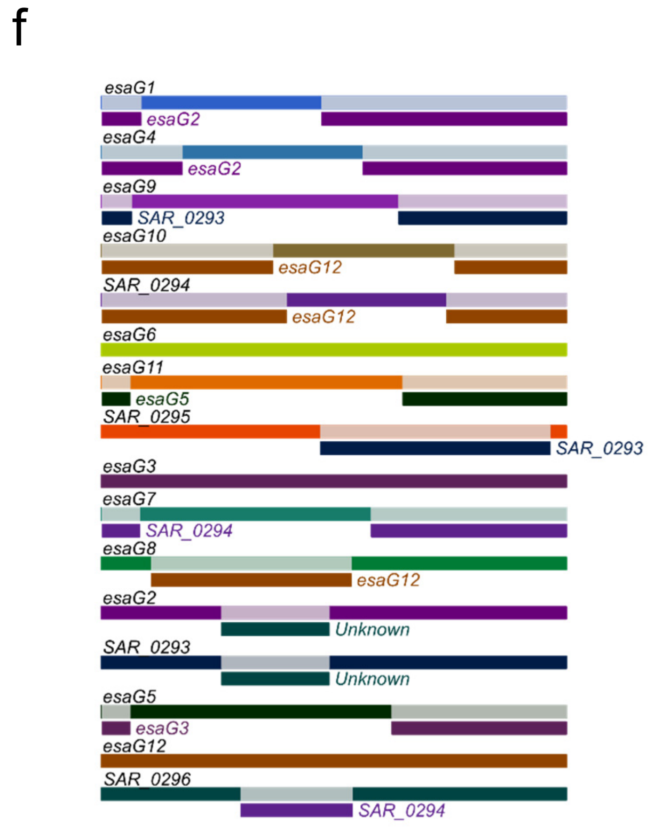


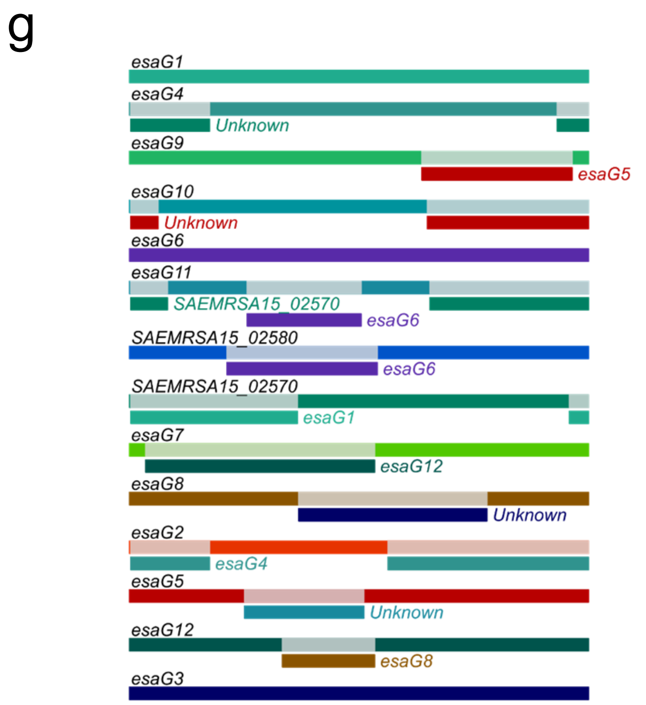


**Fig S4. Recombination events within the *esaG* genes encoded in representative *essC2*, *essC3* and *essC4* variant strains.** a-c. The genes downstream of *essC* differ between the *essC* variants. The regions spanning *essC* to the conserved gene *SAOUHSC_00279* were aligned, using BLAST, between RN6390, and a. ST398 (*essC2* variant), b. MRSA252 (*essC3* variant) and c. HO 5096 0412 (*essC4* variant). Alignments were visualised using genoPlotR, and in this output, regions of homology are highlighted by red-connecting blocks, with the colour intensity of these blocks reflects the percentage identity found between the two compared regions. d. A maximum likelihood tree constructed with IQTREE and annotated in iTOL for all *esaG* homologues found across the four representative *essC* variant strains RN6390, ST398, MRSA252 and HO 5096 0412. e-g. Alignments of *esaG* homologues from RN6390 with e. ST398, f. MRSA252 and g. HO 5096 0412 were analysed using RDP4 to analyse recombination events. Each gene is labelled in black, with regions of recombination labelled directly below in the colour of the gene from which the recombinant section originated. Note that the *esaG* pseudogene in ST398 is covered by the two small genes *SAPIG_0314-0315*, and but here is referred to as a single pseudogene which we annotated as *SAPIG_0314*.

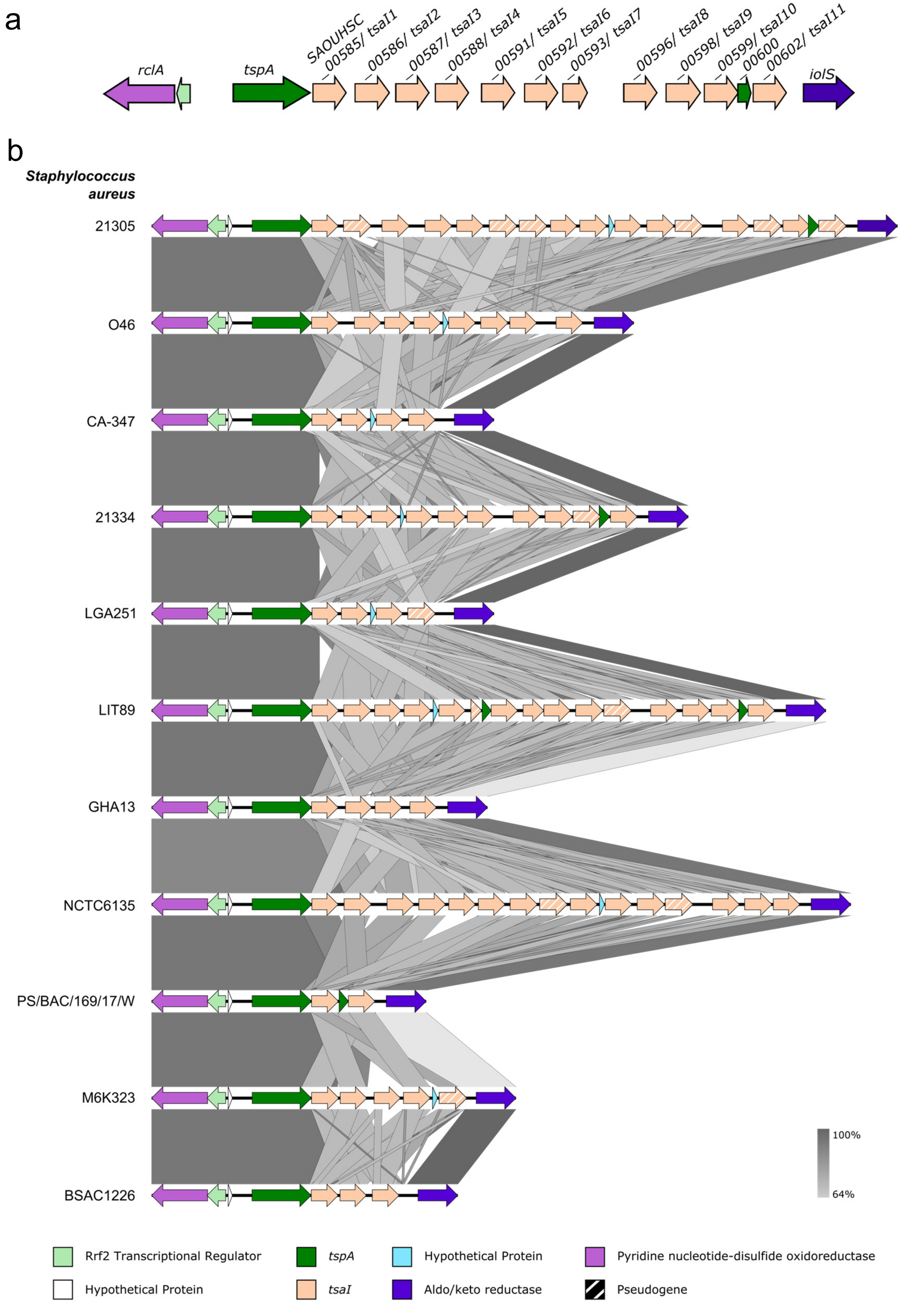
a
rclA
tspA
SAOUHSC
_00585/ tsaI1
_00586/ tsaI2
_00587/ tsaI3
_00588/ tsaI4
_00591/ tsaI5
_00592/ tsaI6
_00593/ tsaI7
_00596/ tsaI8
_00598/ tsaI9
_00599/ tsaI10
_00600
_00602/ tsaI11
iolS
b
Staphylococcus aureus
21305
O46
CA-347
21334
LGA251
LIT89
GHA13
NCTC6135
PS/BAC/169/17/W
M6K323
BSAC1226
100%
64%
Rrf2 Transcriptional Regulator
tspA
Hypothetical Protein
Pyridine nucleotide-disulfide oxidoreductase
Hypothetical Protein
tsaI
Aldo/keto reductase
Pseudogene

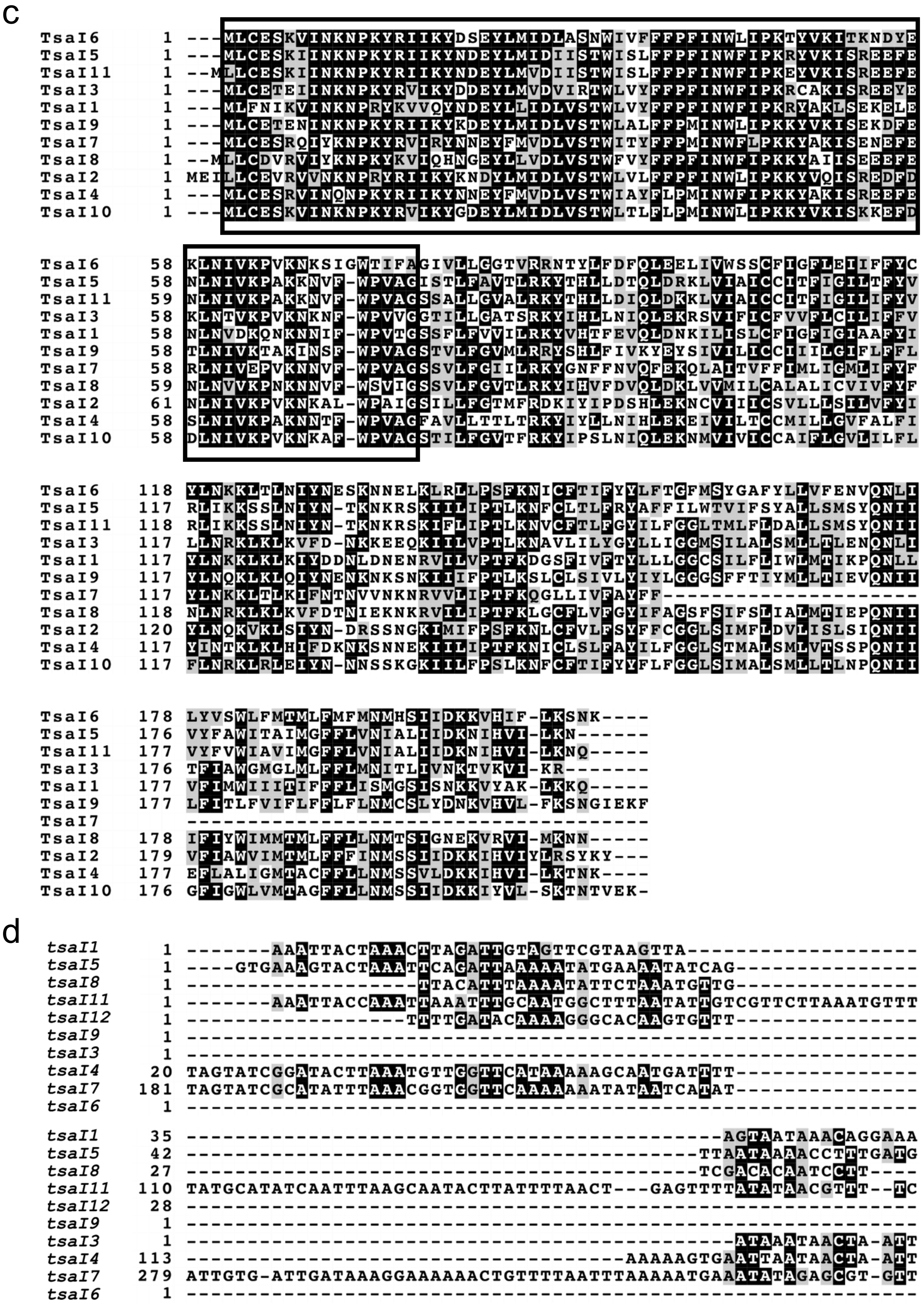


**Fig S5. Multiple *tsaI* homologues are encoded in *S. aureus* strains.** a and b. Genetic arrangement of *tsaI* genes in a. RN6390, and b. a selection of other *S. aureus* strains to demonstrate the variability at this locus. *rclA* and *iolS* are a conserved gene found flanking the TspA locus in *S. aureus* strains, encoding a pyridine nucleotide-disulfide oxidoreductase and an aldo-keto reductase, respectively. c. An alignment of the RN6390 TsaI homologues. The black boxes represent regions of high sequence similarity based in this alignment. d. Alignment of the intergenic region downstream of each RN6390 *tsaI* gene.

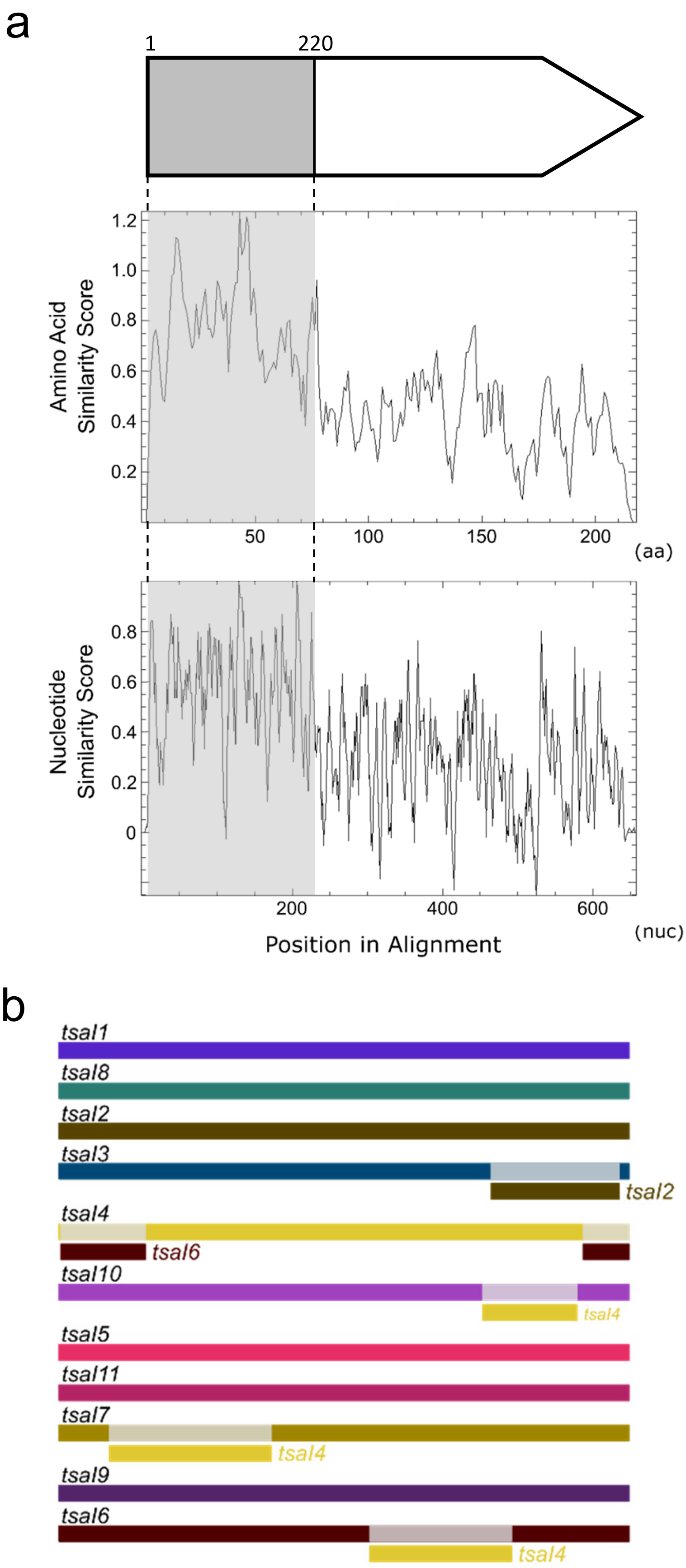


**Fig S6. Assessing recombination events within *tsaI* homologues.** a. A single region of high sequence similarity across the RN6390 TsaI protein sequences (middle panel) and the corresponding nucleotide sequences (bottom panel). The numbers which dictate the limits of the conserved region are taken form the nucleotide sequence of *tsaI1*. b. RDP4 was used to predict recombination events within the *tsaI* homologues encoded in RN6390. Each gene is labelled in black, with regions of recombination labelled directly below in the colour of the gene from which the recombinant section originated.

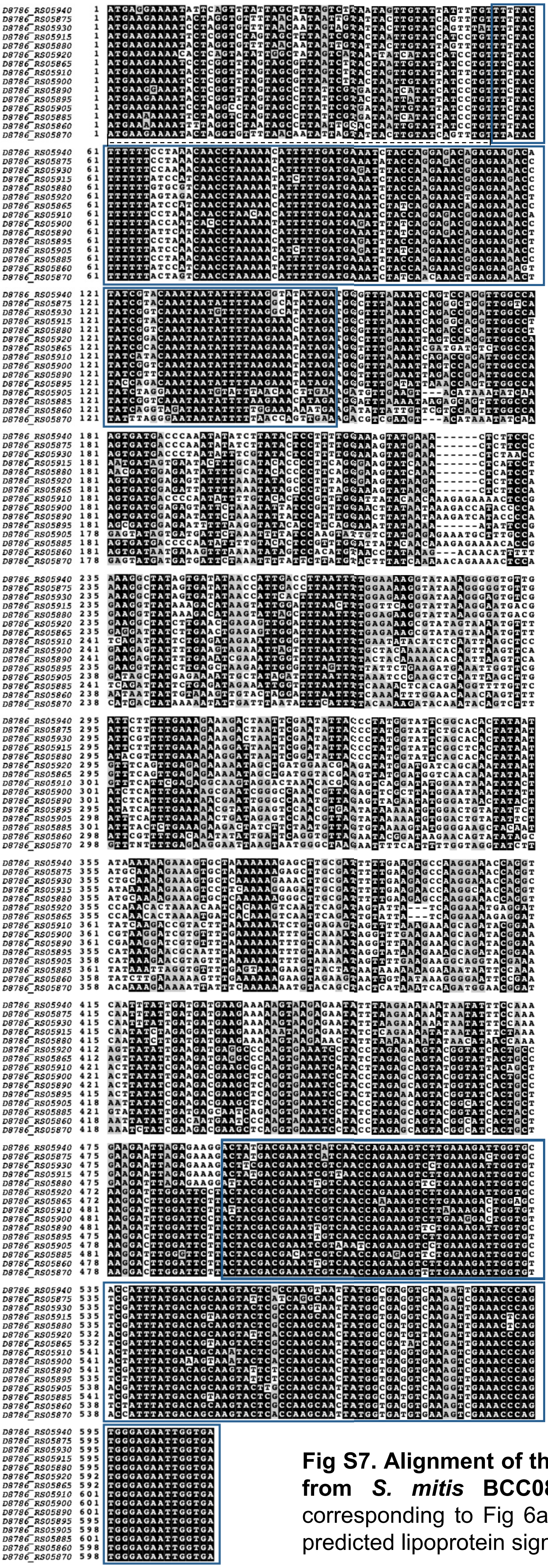

**Fig S7. Alignment of the nucleotide sequences of *tipC* homologues from *S. mitis* BCC08.** The blocks of high sequence similarity corresponding to Fig 6a are outlined in blue. The region encoding the predicted lipoprotein signal peptide is indicated with a dashed line.
